# Protease Resistance of *ex vivo* Amyloid Fibrils implies the proteolytic Selection of disease-associated Fibril Morphologies

**DOI:** 10.1101/2021.07.05.451219

**Authors:** Jonathan Schönfelder, Peter Benedikt Pfeiffer, Tejaswini Pradhan, Johan Bijzet, Bouke P.C. Hazenberg, Stefan O. Schönland, Ute Hegenbart, Bernd Reif, Christian Haupt, Marcus Fändrich

## Abstract

Several studies recently showed that *ex vivo* fibrils from patient or animal tissue were structurally different from *in vitro* formed fibrils from the same polypeptide chain. Analysis of serum amyloid A (SAA) and Aβ-derived amyloid fibrils additionally revealed that *ex vivo* fibrils were more protease stable than *in vitro* fibrils. These observations gave rise to the proteolytic selection hypothesis that suggested that disease-associated amyloid fibrils were selected inside the body by their ability to resist endogenous clearance mechanisms. We here show, for more than twenty different fibril samples, that *ex vivo* fibrils are more protease stable than *in vitro* fibrils. These data support the idea of a proteolytic selection of pathogenic amyloid fibril morphologies and help to explain why only few amino acid sequences lead to amyloid diseases, although many, if not all, polypeptide chains can form amyloid fibrils *in vitro*.

## Introduction

Amyloid fibrils are pathogenic agents in a range of debilitating diseases from systemic amyloidosis to neurodegeneration [1]. They consist of endogenous polypeptide chains that are assembled into cross-β sheets, as demonstrated by X-ray diffraction or other techniques [2]. Amyloid fibrils show conserved structural features, such as a width of ∼10 nm and affinity for Congo red and thioflavin T dyes [3,4]. Around 35 non-homologous human polypeptide chains give rise to amyloid fibrils in the course of disease [5], for example immunoglobulin light chains (LCs) in systemic AL amyloidosis [6], serum amyloid A (SAA) 1.1 protein in systemic AA amyloidosis [7] or transthyretin (TTR) in systemic ATTR amyloidosis [8].

These proteins form fibrils *in vitro* that reproduce the linear transmission electron microscopy (TEM) morphology, dye binding and X-ray diffraction characteristics of amyloid fibrils extracted from diseased tissue [9–11]. The Nomenclature Committee of the International Society of Amyloidosis (ISA) nevertheless recommended for many years that *in vitro* formed cross-β fibrils should be called “amyloid-like” as it was felt that their relationship to *bona fide* amyloid fibrils was not sufficiently established [12]. Indeed, more and more studies recently demonstrated that *in vitro* formed fibrils are structurally different from amyloid fibrils that were purified from amyloidotic tissue, termed here *ex vivo* fibrils. Examples hereof include the fibrils from LC [13,14], tau protein [15], Aβ peptide [16], TTR [17], SAA1.1 [18], α-synuclein [19] and prion protein fibrils [20]. Yet, ISA dropped the distinction between amyloid and amyloid-like fibrils in 2018 [21].

An interesting feature of amyloid fibrils from different patients and animals is that these are structurally conserved, as long as the respective patients or animals are affected by the same disease variant and express the same allele of the fibril protein [13,16,17,19,22–24]. This situation contrasts to *in vitro* fibrils, where substantially different structures may be formed from the same polypeptide chain in different laboratories or under different conditions of fibril formation [25,26]. *Ex vivo* amyloid fibrils from murine SAA1.1 protein or Aβ peptide were additionally found to be more protease stable than their *in vitro* formed counterparts [16,18]. These observations gave rise to the proteolytic selection hypothesis, which assumes that disease-associated fibril morphologies were selected inside the body by their ability to escape endogenous proteolytic clearance systems [18].

We here test this hypothesis by comparing the proteolytic stability of a range of different *ex vivo* and *in vitro* formed amyloid fibrils. We find that all tested samples of *ex vivo* amyloid fibrils are significantly resistant to proteolytic digestion, while all, but one, samples of *in vitro* formed fibrils are readily degraded.

## Materials and Methods

### Extraction of amyloid fibrils from diseased tissue

Amyloid fibrils were extracted from human heart (human AL and ATTR amyloidosis) or kidney (systemic AA amyloidosis) or from murine liver (murine AA amyloidosis) by using a modified version of a previously described extraction protocol [22]. For further details see Supplementary Information.

### TEM analysis of the fibril samples

TEM specimens were prepared by placing 3 µL of a fibril protein solution onto a formvar and carbon coated 200 mesh copper grid (Plano) which was freshly glow discharged with a PELCO easiGlow instrument (TED PELLA). The sample was incubated on the grid for 3 min before it was soaked off with filter paper (Whatman). The grid was washed three times with 7 µL pure water, each time followed by the removal of the water with the filter paper. The specimen was stained briefly with 7 µL of a 2 % (w/v) uranyl acetate solution, which was removed with filter paper. The staining procedure was repeated two more times. The grids were dried for at least 5 min before they were examined in a JEM-1400 TEM (JEOL), operated at 120 kV and equipped with a F216 camera (TVIPS). All pictures were taken at a magnification of 20.000 x.

### Protease digestion of fibrils

Amyloid fibrils were subjected to a protease digestion protocol to assess their proteolytic stability. For proteinase K digestion, we mixed fibril stocks with pure water and 22 µL of 10x TCB buffer [200 mM Tris, 1.4 M NaCl, 20 mM CaCl2, 1 % (w/v) NaN3, pH 8.0] to achieve a final volume of 220 µL with a fibril protein concentration of 200 or 100 µg/mL, respectively. The protein concentration in the stocks of the *ex vivo* fibrils were determined as will be described in the next section. The protein concentrations in the *in vitro* fibril stocks were used as they were prepared. The reaction tube was transferred to a heating chamber (FED 115 APT.Line, Binder) at 37 °C and the digestion was started by addition of 0.4 μL proteinase K (20 mg/mL, Thermo Fisher Scientific). For the digests of fibrils in the lower concentration of 100 µg/mL only 0.2 µL of proteinase K were used. A first aliquot (20 µL) was removed immediately prior to the addition of proteinase K. All further aliquots were removed 5, 10, 30 and 60 min after the protease had been added to the reaction tube. As soon as an aliquot was taken, 2 µL of a protease inhibitor solution [200 mM phenylmethylsulfonyl fluoride (PMSF) in methanol, Carl Roth] were added and the mixture was incubated for 10 min at room temperature and frozen in liquid nitrogen. Once all aliquots were collected, they were thawed at room temperature and prepared for denaturing gel electrophoresis.

For pronase E (Sigma) digests we used essentially the same procedure, except that all digests contained 200 µg/mL fibrils and 40 µg/mL pronase E and that the digest was stopped in each 20 µL aliquot with 2 µL protease inhibitor cocktail solution (1 tablet cOmplete EDTA-free Protease Inhibitor Cocktail, Roche, in 2 ml pure water) instead of PMSF solution. The pronase E solution was freshly prepared each time by dissolving 20 mg pronase E in 1 mL pure water.

### Determination of the fibril protein concentration in the ex vivo fibril stocks

The concentration of the fibril proteins were determined by denaturing gel electrophoresis by comparison with a protein standard, which was prepared from recombinant murine SAA1.1 protein. 1 mg murine SAA1.1 protein were dissolved in 1 ml pure water. A 200 μL aliquot of this solution was mixed with 800 μL 7.5 M guanidine hydrochloride in 25 mM sodium phosphate buffer, pH 6.5. Based on the extinction of this solution at 280 nm, which was measured with a Lambda Bio+ spectrometer (Perkin Elmer), and the theoretic molar extinction coefficient of 24,750 [M^−1^ cm^−1^] for murine SAA1.1 protein the concentration was determined according to the Lambert-Beer law. This SAA1.1 solution was used to prepare a protein standard series with protein concentrations ranging from 50 to 500 μg/mL. The standard series was run side-by-side with the ex vivo fibril stocks on a denaturing protein gel, which was densitometrically analysed with ImageJ [27]. The band intensity of the standard (n=2) was plotted as a function of standard protein concentration.

### Denaturing protein gel electrophoresis

To analyze the proteolytic digest, each aliquot (22 µL) was mixed with 7,3 µL 4× NuPAGE LDS Sample Buffer (Thermo Fisher Scientific) by repeated pipetting. The samples were heated at 95 °C for 10 min in a block heater. A gel chamber with a 26 well 4−12% NuPAGE Bis-Tris Gels (Thermo Fisher Scientific) was assembled and filled with 1X NuPAGE MES-SDS Running Buffer (Thermo Fisher Scientific). A total of 10 µL per sample were loaded onto the gel, together with 8 µL of BlueEasy prestained protein marker (Nippon Genetics GmbH). All gels were run for 35 min at 180 V and at room temperature and stained in Coomassie staining solution [2.5 % (w/v) Coomassie Brilliant Blue R250, 30 % (v/v) ethanol, 10 % (v/v) acetic acid] for 1 h at room temperature on a platform shaker (Polymax 1040, Heidolph). Hereafter, the gels were transferred into destaining solution [20 % (v/v) ethanol, 10 % (v/v) acetic acid] and incubated for 20 h. Gel images were taken by using the Box Chemi XL1.4 (Syngene G).

## Results

### Ex vivo *amyloid fibrils are highly proteinase K resistant*

We analyzed *ex vivo* amyloid fibrils from three major forms of systemic amyloidosis (AA, AL and ATTR). AA amyloid fibrils were extracted from a human patient suffering from the common/glomerular disease variant and from two AA amyloidotic mice. The fibrils are derived from SAA1.1 protein in both species. ATTR amyloid fibrils were purified from three patients that were heterozygous for the TTR mutations Gly47Asp (patient ATTR-H03) and Val20Ile (patients ATTR-H04 and ATTR-H14). The three ATTR cases showed type A ATTR fibrils (Fig. 1); that is, their fibrils consist of full-length TTR as well as TTR fragments [28]. AL amyloid fibrils were purified from explanted hearts of patients FOR001, FOR005 [14] and FOR006 [29] with λ-AL amyloidosis and severe cardiomyopathy. Cases FOR001 and FOR006 are associated with a λ1 LC, FOR006 with a λ3 LC.

**Figure 1.**
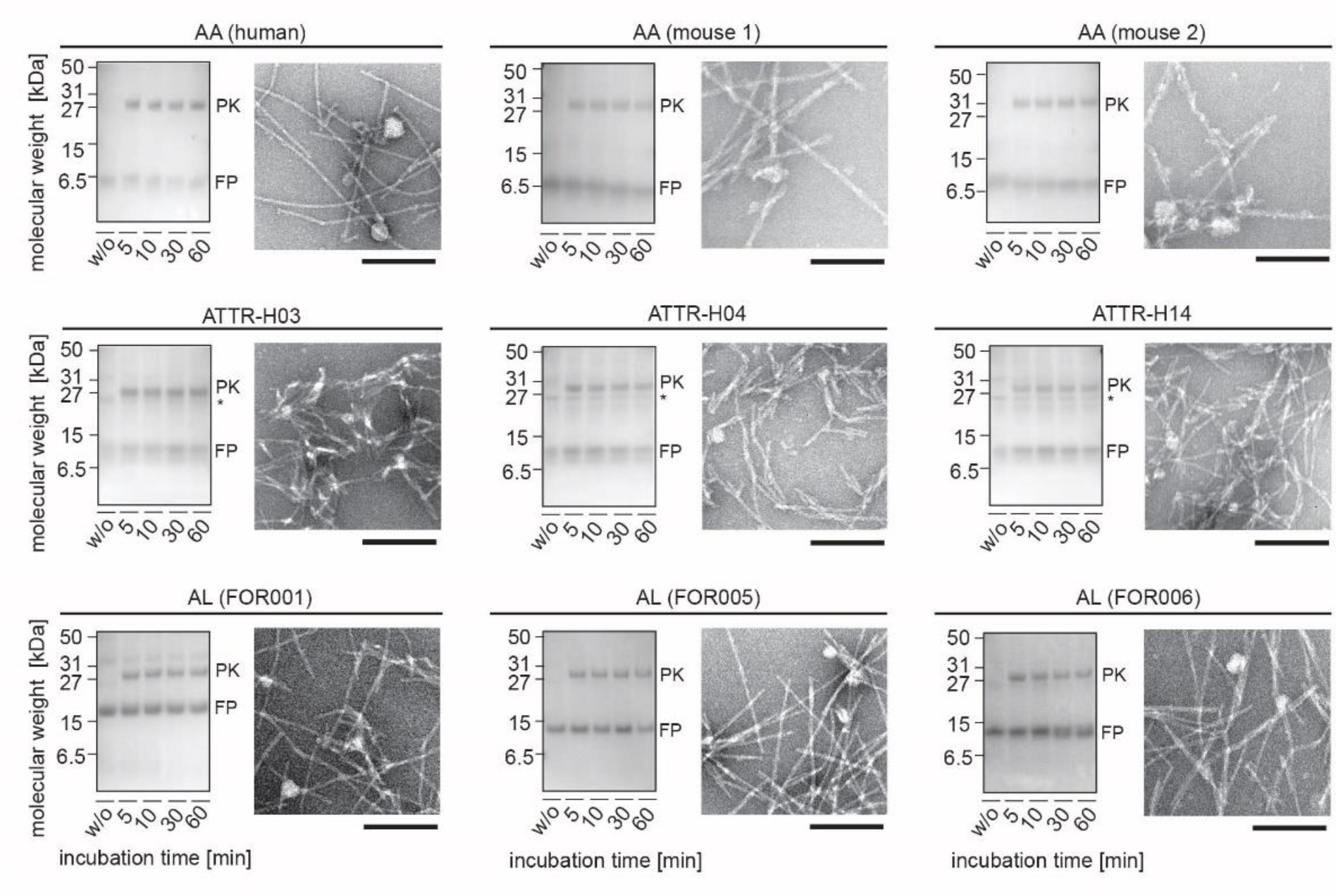
*Ex vivo* amyloid fibrils are highly proteinase K stable. Nine cases of *ex vivo* amyloid fibrils were examined. The left part of each panel shows a Coomassie stained denaturing protein gel of the fibrils digested with proteinase K for 5, 10, 30 or 60 min, as indicated in the Figure. The first lane shows fibril sample before proteinase K addition. FP: fibril protein; PK: proteinase K. *: SAP. The samples contained a nominal fibril protein concentration of 200 µg/mL and 40 µg/mL proteinase K. Gels of digests using a nominal fibril protein concentration of 100 µg/mL and 20 µg/mL proteinase K are shown in Fig. S1. The right part shows a TEM image of the undigested fibrils. Scale bar: 200 nm.

TEM confirmed, for all samples, the linear morphology of the extracted filaments and their abundance in negatively stained TEM specimens (Fig. 1). The fibrils were exposed to proteinase K digestion and aliquots were removed at different time points (5, 10, 30 and 60 min after addition of the protease). Proteolysis was stopped with PMSF and the samples were analyzed with denaturing polyacrylamide gel electrophoresis. All fibril proteins were stable under the harsh proteolytic conditions of our experiment and persisted until the end of the experiment (Fig. 1). There was no major difference between the different types of systemic amyloidosis or between humans and murine samples. Essentially the same result was obtained with a second series of samples in which we reduced the fibril and protease concentrations by 50 % (Fig. S1). All tested samples of *ex vivo* amyloid fibrils presented significant resistance to protease digestion.

### In vitro *fibrils from disease-associated polypeptide chains are proteinase K sensitive*

We prepared amyloid fibrils *in vitro* from chemically synthetic, recombinantly expressed or commercially available polypeptide chains (SI Table 1). We first focused on amyloid fibrils from polypeptide chains, which, according to the definitions of ISA, give rise to amyloid diseases [5]: human and murine SAA1.1, TTR, a recombinant immunoglobulin LC fragment corresponding to the FOR005 variable LC domain, human lysozyme, Aβ(1-40) peptide, α-synuclein and human insulin. We refer to this set of fibrils as ‘*in vitro* fibrils from disease-associated polypeptide chains’. Using TEM the samples are rich in amyloid fibrils, although the fibrils differed in morphology. Some fibrils were straight and thick, such as murine SAA1.1 and Aβ(1-40) fibrils, while others, such as human SAA1.1 and TTR fibrils, were relatively thin and curvilinear (Fig. 2).

**Figure 2.**
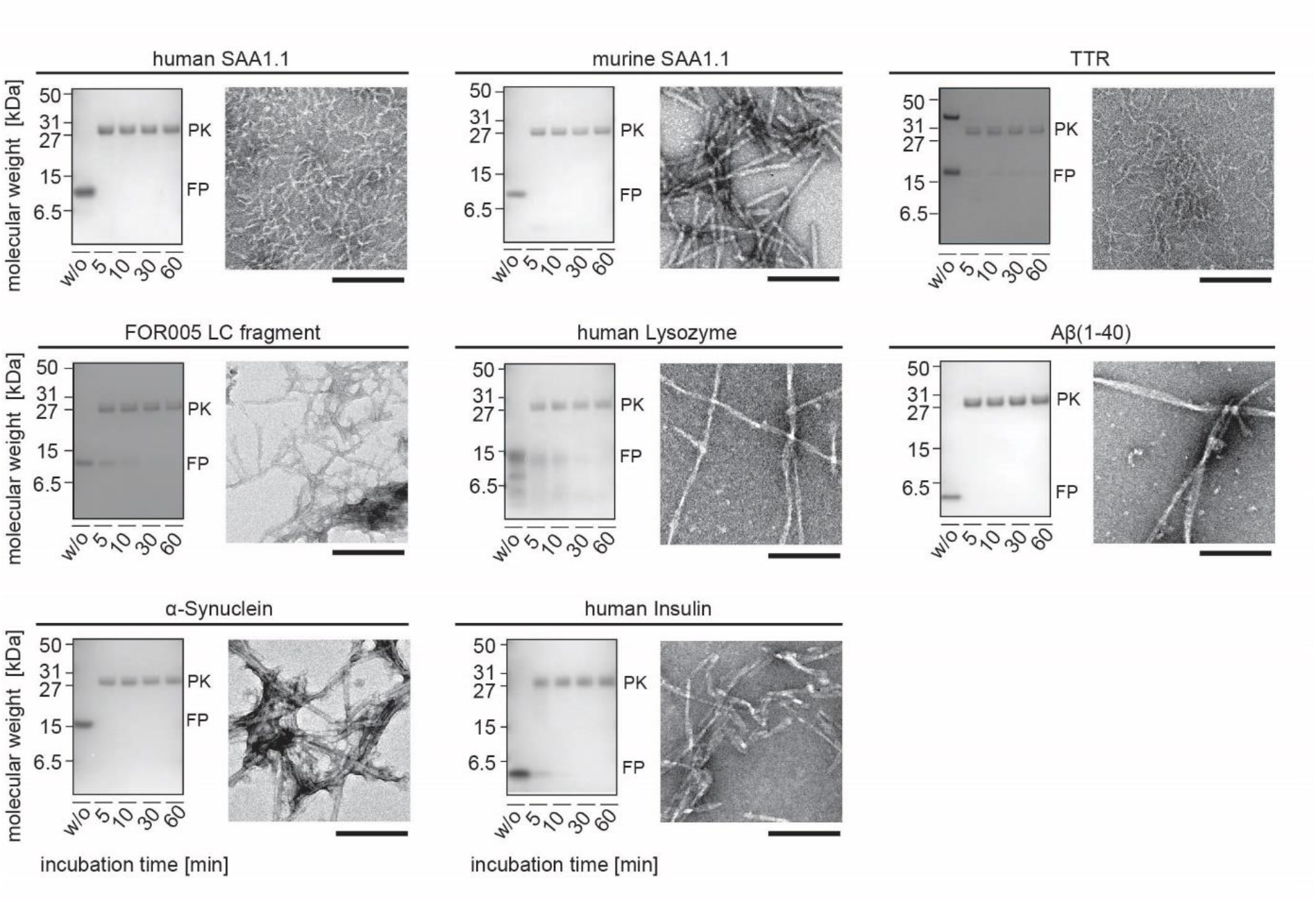
*In vitro* fibrils from disease-associated polypeptide chains are proteinase K sensitive. Eight cases of *in vitro* fibrils from disease-associated polypeptide chains were examined. The left part of each panel shows a Coomassie stained denaturing protein gel of the fibrils digested with proteinase K for 5, 10, 30 or 60 min, as indicated in the Figure. The first lane shows fibril sample before proteinase K addition. FP: fibril protein; PK: proteinase K. The samples contained a nominal fibril protein concentration of 200 µg/mL and 40 µg/mL proteinase K. Gels of digests using a nominal fibril protein concentration of 100 µg/mL and 20 µg/mL proteinase K are shown in Fig. S2. The right part shows a TEM image of the undigested fibrils. Scale bar: 200 nm.

We then subjected the fibrils to the same proteolytic conditions, which we used above to analyze the proteolytic stability of *ex vivo* amyloid fibrils (Fig. 1, Fig. S1). Remarkably, all analyzed samples of *in vitro* fibrils from disease-associated polypeptide chains were readily degraded. In most cases we find the fibril proteins to become fully destroyed within 5 min (Fig. 2). Similar to *ex vivo* fibrils, we obtained consistent results if we reduced the fibril and protease concentrations in the reaction vessels by 50 % (Fig. S2). *In vitro* fibrils from disease-associated polypeptide chains are less stable to proteolysis than *ex vivo* amyloid fibrils, at least under the presently used experimental conditions.

### *Most* in vitro *fibrils from non-pathogenic polypeptide chains are proteinase K sensitive*

In a next step, we analyzed *in vitro* formed amyloid fibrils from polypeptide chains that are not related to amyloid diseases. Some of the polypeptide chains are currently under investigation as to whether they form amyloid deposits *in vivo*, such as glucagon [5]. Others are not suspected at all to form amyloid fibrils *in vivo*. We refer to this set of fibrils as ‘*in vitro* fibrils from non-pathogenic polypeptide chains’. They were derived from glucagon, apomyoglobin, bovine insulin, α-lactalbumin, β-lactoglobulin, α-crystallin and the 248-286 residue fragment of prostatic acidic phosphatase, that is commonly referred to as SEVI (‘semen derived enhancer of viral infection’) [30]. Subjected to proteinase K digestion, we find that the majority of *in vitro* fibrils from non-pathogenic polypeptide chains are readily degraded (Fig. 3, Fig. S3). The only exception is sample of α-lactalbumin fibrils that were stable until the end of the experiment (60 min), although the intensity of the α-lactalbumin band become progressively weaker the longer these fibrils were exposed to proteinase K (Fig. 3, Fig. S3). We conclude that *in vitro* fibrils from non-pathogenic polypeptide chains are not very stable to proteinase K digestion, although our results with α-lactalbumin show that proteolytic resistant fibrils can be formed to some extent within a test tube.

**Figure 3.**
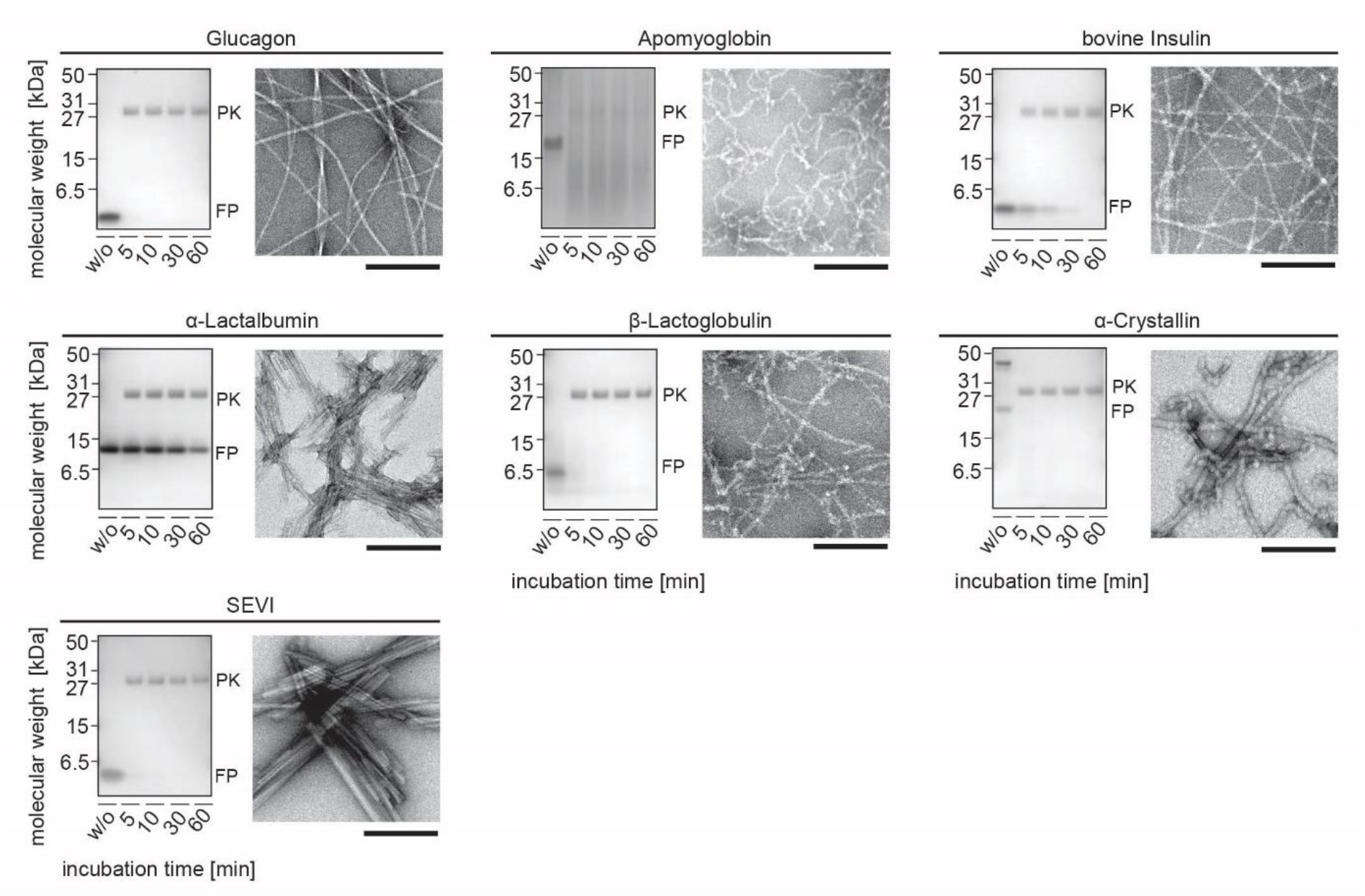
Most *in vitro* fibrils from non-pathogenic polypeptide chains are proteinase K sensitive. Seven cases of *in vitro* fibrils from non-pathogenic polypeptide chains were examined. The left part of each panel shows a Coomassie stained denaturing protein gel of the fibrils digested with proteinase K for 5, 10, 30 or 60 min, as indicated in the Figure. FP: fibril protein; PK: proteinase K. The samples contained a nominal fibril protein concentration of 200 µg/mL and 40 µg/mL proteinase K. Gels of digests using a nominal fibril protein concentration of 100 µg/mL and 20 µg/mL proteinase K are shown in Fig. S3. The first lane shows fibril sample before proteinase K addition. The right part shows a TEM image of the undigested fibrils. Scale bar: 200 nm.

### Ex vivo *fibrils are also more pronase E stable than most* in vitro *fibrils*

Since all above findings are based on a single protease (proteinase K), we additionally tested several samples with pronase E. We previously found with *ex vivo* and *in vitro* fibrils from murine SAA1.1 protein that *ex vivo* fibrils are more protease stable than *in vitro* fibrils irrespective of whether we used proteinase K, pronase E, leucine aminopeptidase or carboxypeptidease A [18]. Pronase E is a mixture of different proteases from *Streptomyces griseus* [31]. We find that all samples of *ex vivo* amyloid fibrils (human AA, murine AA and human ATTRG47D patient ATTR-H03) are highly pronase E stable (Fig. 4), while the majority of the tested *in vitro* formed fibrils (human SAA1.1, α-synuclein, SEVI, α-crystallin and glucagon) become rapidly degraded under these conditions (Fig. 4). Only the sample containing α-lactalbumin fibrils was resistant to digestion with pronase E (Fig. 4), corroborating our observations with proteinase K (Fig. 3, Fig. S3). In conclusion, *in vitro* fibrils are usually more protease sensitive than *ex vivo* fibrils.

**Figure 4.**
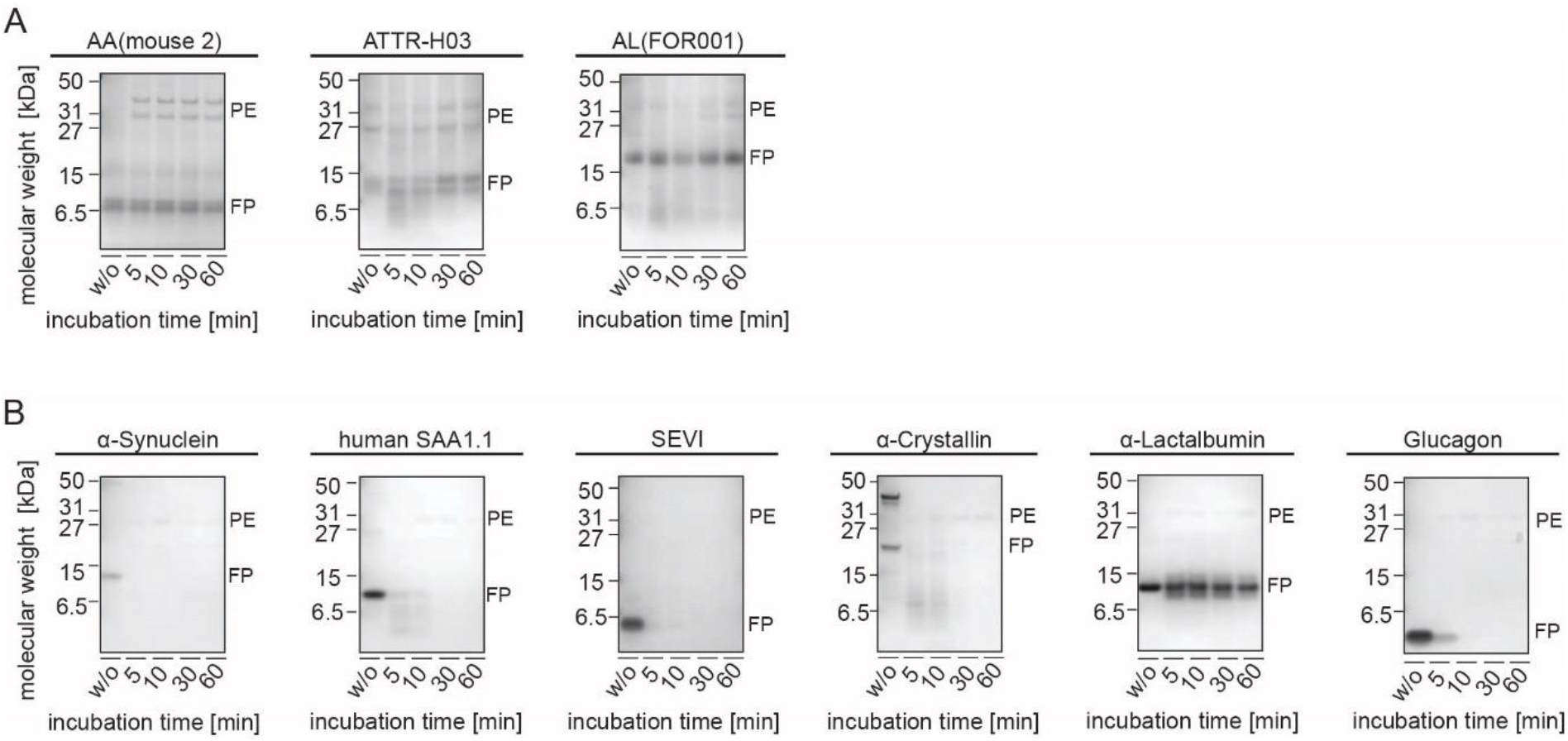
*Ex vivo* fibrils are more pronase E stable than most *in vitro* fibrils. (A) Coomassie stained denaturing protein gel of the *ex vivo* fibrils digested with pronase E for 5, 10, 30 or 60 min, as indicated in the Figure. FP: fibril protein; PE: pronase E. (B) Coomassie stained denaturing protein gels of the *in vitro* fibrils treated accordingly.

### *The observed proteolytic stability of* ex vivo *amyloid does not arise from SAP*

To interrogate the possible molecular basis of this effect we analyzed the possible contributions of serum amyloid P component (SAP). Similar to previous analyses which showed that *ex vivo* fibrils can contain SAP [17], we find significant amounts of SAP in our *ex vivo* fibrils, and specifically in ATTR amyloid fibrils (Fig. 1). Western Blot-based quantifications of SAP in samples containing 200 μg/mL fibril protein revealed SAP concentration of 38.2 ± 1.5 μg/mL for the three samples of ATTR amyloid fibrils and of 3.4 ± 2.3 μg/mL for the four samples of human AA and AL amyloidosis (Fig. S4).

SAP represents an important non-fibril component of amyloid deposits, that protects these deposits from being cleared *in vivo* [32]. It also counteracts the degradation of amyloid fibrils *in vitro* [33], suggesting that the presently described differences between (SAP-free) *in vitro* and (SAP-containing) *ex vivo* fibrils might have originated from SAP. Therefore, we spiked our samples of *in vitro* fibrils with SAP, which we had purified form the amyloid tissue of a patient with systemic ATTR amyloidosis. We investigated *in vitro* formed fibrils from human SAA1.1, α-synuclein, SEVI, α-crystallin and glucagon (Fig. S4). Some samples, such as the ones prepared from SEVI or glucagon, show a slightly retarded proteolysis compared with SAP-free samples (Fig. 2) that is consistent with a reduced proteolysis in the presence of SAP. However, none of the SAP-containing samples showed a proteolytic stability that resembled remotely the proteolytic resistance of the *ex vivo* amyloid fibrils seen in the experiments reported in Fig. 1 and Fig. S1. We conclude that the simple presence of SAP cannot explain the observed proteolytic stability of *ex vivo* amyloid fibrils.

## Discussion

Several studies previously showed that disease-associated amyloid fibrils from patient tissue are structurally different from fibrils formed from the same polypeptide chain *in vitro* [13–16,18–20]. In two cases, namely murine SAA1.1 protein and human Aβ peptide, it was additionally found that *ex vivo* fibrils are more protease resistant than *in vitro* fibrils [16,18]. These observations led to the hypothesis that disease-associated amyloid fibrils were selected inside the body by their proteolytic stability [18]. That is, protease stable fibrils have a higher chance to escape endogenous clearance mechanisms, to proliferate and to accumulate in a native cellular environment. This mechanism was termed proteolytic selection [18]. Our new observations further support this hypothesis by showing, for a range of amyloid fibrils, that *ex vivo* fibrils are more stable to proteolysis than *in vitro* fibrils.

That amyloid fibrils or prions are more resistant to proteolysis than natively folded, globular proteins is known for many years [34–36]. Our study demonstrates, in addition, differences in the stability of fibrils from different sources and that *ex vivo* amyloid fibrils are more stable to proteolysis than most *in vitro* fibrils. Although our results are mainly based on mostly extracellular *ex vivo* amyloid fibrils, such as AA, AL, ATTR amyloid fibrils, and Aβ peptide fibrils from Alzheimer’s disease and cerebral amyloid angiopathy [16], deposits from intracellular amyloid proteins were also described as relatively protease stable, for example deposits from α-synuclein [37,38] or tau protein [39,40].

The observed higher proteolytic stability of *ex vivo* fibrils does not mean that it is impossible to form fibrils *in vitro* that are highly protease stable. Our data with α-lactalbumin show, in fact, that this can be achieved (Fig. 3, Fig. 4; Fig. S3). However, the fibril morphology is known to depend on the conditions of fibril formation [25,26], and we would therefore expect that a polypeptide chain, such as SAA1.1 or TTR, will also be able to form highly protease-stable fibrils inside a test tube, provided that the conditions are right. Yet, it is currently unknown what a ‘right condition’ really is, and it is even possible that differences exist for sequentially different polypeptide chains.

Two possible reasons can be provided as to why fibrils, which were extracted from diseased tissue, are more proteolytically stable than *in vitro* fibrils. The first one attributes encountered differences in proteolytic susceptibility to differences in the fibril structure, which is supported by studies revealing structural differences between *ex vivo* and *in vitro* fibrils [13–16,18–20]. The different fibril structures may be associated with different thermodynamic stabilities of the fibrils, although protease resistant mouse AA amyloid fibrils were recently found not to be very stable to guanidine denaturation [18]. The second one is associated with the fact that *ex vivo* amyloid fibrils can contain non-fibril components of amyloid tissue deposits, such as SAP, which was previously found to protect amyloid fibrils from being degraded *in vitro* [33] and *in vivo* [32]. We find that our *ex vivo* fibrils contain significant amounts of SAP (Fig. S4), but the weight ratios differed for different types of amyloidosis, and much more SAP could be co-purified with ATTR amyloid fibrils than with AA or AL amyloid fibrils (Fig. S4).

When the effect of SAP was tested in our proteolytic assay at a 1:10 weight ratio (SAP to fibril protein), which is within the range of weight ratios in our fibril extracts (Fig. S4), we find only a modest, if any, increase in the proteolytic resistance (Fig. 5). These results are in accord to previous data that a 40-50 % reduction in the digestion of *in vitro* Aβ fibrils with pronase was only observed at a weight ratio of 1:1 (SAP to aged Aβ), while a 50-80% reduction required at a weight ratio of 10:1 (SAP to aged Aβ) [33]. These weight rations are significantly higher than the ones seen in our fibril extracts (Fig. S4), demonstrating that the simple presence of SAP cannot explain the stark differences in proteolytic stability of *ex vivo* and *in vitro* fibrils observed here. This conclusion is corroborated by the fact that one of our samples, namely the one prepared from α-lactalbumin, is highly protease stable in the absence of any SAP (Fig. 3, Fig. 4, Fig. S3). However, amyloid deposits may contain non-fibril components beyond, such as glycosaminoglycans [41], lipids [42], vitronectin or apolipoprotein E [43]. Up to 12 % (w/w) of the anhydrous mass of ex vivo fibrils were reported to be originate from lipids, with sphingolipids, cholesterol and cholesterol esters being in particular abundant [42]. Up to 2 % (w/w) of the mass can be glycosaminoglycans [44], in particular heparan sulfate and dermatan sulfate [45]. We cannot rule out that SAP in conjunction with other non-fibril components of amyloid deposits that were not tested in our study contribute to defining the proteolytic stability of *ex vivo* amyloid fibrils.

**Figure 5.**
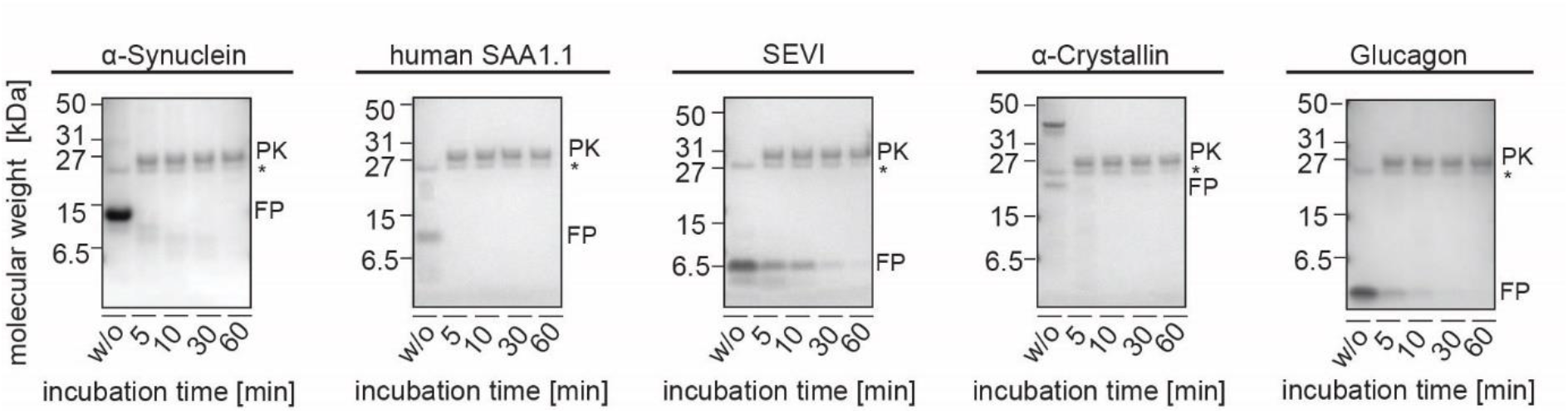
SAP does not substantially protect amyloid fibrils from proteolysis *in vitro*. Coomassie stained denaturing protein gel of the fibrils supplemented with SAP that were digested with proteinase K for 5, 10, 30 or 60 min, as indicated in the Figure. FP: fibril protein; PK: proteinase K; *: SAP. The samples contained a nominal fibril protein concentration of 200 µg/mL and 40 µg/mL proteinase K. Prior to addition of the protease, the fibrils were incubated with 20 µg/mL SAP for 24 h at 37°C under constant agitation (300 rpm).

The present observations are finally relevant in the context of a long-standing debate regarding the molecular basis of amyloid fibril diseases. The work of the late Chris Dobson and his coworkers suggested that amyloid fibrils represent a generic structural state of polypeptide chains and that many, if not all, amino acid sequences are able to form amyloid fibrils [46–48]. An obvious question that was raised by these ideas is: why are only ∼35 non-homologous protein sequences associated with amyloid diseases *in vivo* [5], although the human body contains thousands of non-homologous polypeptide chains? At least two factors could be put forward previously to explain this paradox.

First, efficient folding into a globular protein counteracts amyloid formation, as shown for example for systemic ATTR amyloidosis [49] and lysozyme misfolding in systemic Alys amyloidosis [50]. Second, amyloid fibril formation requires high concentrations of the fibril precursor protein, and many classical fibril precursor proteins are abundant inside the body, at least in certain medical situations. The need of a high protein concentration is illustrated by SAA1.1 and LCs. SAA1.1 is an acute phase protein that is strongly upregulated during inflammation, and chronic inflammatory conditions are a precondition to development of systemic AA amyloidosis [7]. Similarly, a highly concentrated monoclonal free LC in the serum, that arises from a plasma cell dyscrasia, is typically the precondition to fibril formation in systemic AL amyloidosis [51].

A potential third factor that may be added based on the present findings is the need to form a highly stable fibril state. In the case of native folding reactions that lead to globular proteins it is known that only a limited number of polypeptide chains possesses an appropriate amino acid sequence to form these compact protein states. As a result, there are many so-called intrinsically disordered proteins that are unable to fold into compact globular conformations [52]. In case of amyloid fibrils, we suggest that although the formation of such fibrils is a common property of polypeptide chains with an appropriate backbone structure there will be differences in the stability of fibrils in particular to proteolytic digestion. These differences may arise from the specific packing of fibril protein segments that depend on the amino acid sequence and on the presence of a specific, and complementary, arrangement of side chains. As a result, the number of amino acid sequences that allow the formation of pathogenic amyloid fibrils will be smaller than the number of proteins that can form any type of cross-β fibril.

## Abbreviations

ISA: International Society of Amyloidosis
LC: immunoglobulin light chain
SAA: serum amyloid A
PMSF: phenylmethylsulfonyl fluoride
SAP: serum amyloid P component
TEM: transmission electron microscopy
TTR: transthyretin

## Acknowledgements

The authors thank the tissue donors and their families for enabling this research. We thank K. Danzer, J. W. Kelly and J. Münch for gifts of proteins or peptides.

## Disclosure of interest

The author(s) declare no conflict of interest.

## Funding

This study was supported by the Deutsche Forschungsgemeinschaft, including the support for the Research Unit FOR 2969 (grant numbers FA 456/27, HA 7138/3, HE 8472/1, SCHO 1364/2 and RE1435/19-1) to M.F., C.H., U.H., S.O.S. and B.R. and FA 456/28 to M.F.

## Notes

### Competing Interest Statement

The authors have declared no competing interest.

