## Supplementary Material for "Protease Resistance of *ex vivo* Amyloid Fibrils implies the proteolytic Selection of disease-associated Fibril Morphologies"

### SI Methods

#### *Description of the human amyloidosis samples*

Systemic AA amyloidosis: Kidney tissue was derived from a female patient who suffered from proteinuria and advanced loss of renal function due to AA amyloidosis. The woman died at the age of 70 and the clinical history has been described elsewhere, reported as case AA4 [1]. The cryo-EM structure of the fibrils from this patient has been determined [1]. The kidney tissue was collected at the University Hospital Groningen during autopsy and stored frozen until further analysis.

Systemic ATTR amyloidosis: Heart tissue of three patients with hereditary ATTR amyloidosis has been used to extract ATTR amyloid fibrils. Patient ATTR-H03 was a 62-year-old woman, who carried a heterozygous TTRG47D mutation. Hereditary ATTR amyloidosis (first time in her family) was diagnosed by cardiac biopsy and genetic analysis. She had also carpal tunnel syndrome. Her cardiac failure worsened and she died 41 days after heart transplantation from sepsis. Patient ATTR-H04 was a 55-year-old man heterozygous for the TTRV20I mutation. He was diagnosed with cardiomyopathy and further examinations (biopsy and genetic testing) revealed ATTR amyloidosis of the cardiac phenotype. He is well 8.5 years after heart transplantation. Patient ATTR-H14 was a 49 years old male who was heterozygous for TTRV20I with the same phenotype as patient ATTR-H04. He had a very advanced heart failure and died 3 days after heart transplantation from multi organ failure. The heart tissue was freshly frozen after explantation at the University Clinic Heidelberg and stored at -80 °C until further use.

Systemic AL amyloidosis: AL amyloid fibrils were extracted from the hearts of three patients suffering from advanced heart failure due to AL amyloidosis. In all three cases, the underlying disease was a λ LC restricted monoclonal gammopathy. The patients were treated within the heart transplant program of the University Hospital Heidelberg. All three patients had isolated cardiac involvement. Patient FOR001 was, at the time point of cardiac surgery, a 51-year-old male who showed λ1 LC deposition. He is alive and well 8 years after the procedure. Patient FOR005 was a 50-year-old female whose deposits are derived from λ3 LCs [2,3]. She died 2.5 years later from disease progression. Patient FOR006 was a 51-year-old female who suffered from λ1 LC deposition [2,3]. She is alive and well 6.5 years after the transplant. The explanted heart tissue was stored at -80 °C.

***Ethics statement***

Informed consent was obtained from the patient and/or the relatives for material collection. Material collection was performed by observing all relevant regulations and legal requirements and based on the ethical approval from the ethical committees of the University Clinics of Heidelberg, Germany, (123/2006) and Groningen, The Netherlands, (201800551). The biochemical extraction and analysis at Ulm University were approved by the ethical committee of Ulm University, Germany (210/13).

***Description of the murine amyloidosis samples***

Fibrils were purified from experimentally induced AA amyloidotic mice. Female 6- to 8-week-old NMRI mice (Charles River Laboratories) received a single 0.1 mL injection of 0.1 mg/mL solution of murine AA amyloid fibrils into the lateral tail vein. Immediately afterwards, mice have been subcutaneously injected with 0.2 mL of a freshly prepared 1 % (w/v) solution of $AgNO_3$ in distilled water. The $AgNO_3$ injection was repeated using 0.1 mL after 7 and 14 days. On day 16, animals were euthanized with $CO_2$ and spleens were removed subsequently. AA fibrils from amyloid-laden mouse spleen were extracted as described elsewhere [1,2]. The animal experiments were approved by the Regierungspräsidium Tübingen, Germany (TVA

1165) and all methods were carried out in accordance with relevant guidelines and regulations. All methods are reported in accordance with ARRIVE guidelines (https://arriveguidelines.org) for the reporting of animal experiments.

***Extraction of amyloid fibrils from diseased tissue***

200 mg of diseased tissue were minced with a scalpel to the point where no individual pieces were discernible and transferred into a reaction tube. Next, 500 µL ice-cold Tris-Calcium Buffer [20 mM Tris, 138 mM NaCl, 2 mM $CaCl_2$, 0.1 % (w/v) $NaN_3$, pH 8.0] were added to the tube and the contents were mixed thoroughly by repeated inversion. The tube was then centrifuged for 5 min at 3,100 × g (4 °C) and the resulting supernatant was collected. This initial wash step was repeated for a total of five times. After completion, the pellet was resuspended in 1 mL collagenase solution [1 tablet cOmplete ethylenediaminetetraacetic acid (EDTA)-free Protease Inhibitor Cocktail, Roche, in 7 ml TCB with 5 mg/ml *Clostridium histolyticum* collagenase, Sigma] that had been prepared on the same day and incubated overnight on a MTS 2/4 digital microtiter shaker at 750 rpm (37 °C). Hereafter, the contents of the tube were centrifuged for 30 min at 3,100 × g (4 °C). The resulting supernatant was collected while the pellet was resuspended in 500 µL ice-cold Tris–EDTA buffer [20 mm Tris, 140 mm NaCl, 10 mm EDTA, 0.1 % (w/v) $NaN_3$, pH 8.0] immediately followed by centrifugation for 5 min at 3,100 × g (4 °C). The latter wash step was repeated for a total of five times, and after each centrifugation step the supernatant was collected. To extract the fibrils, the pellet was ultimately resuspended in 500 µL ice-cold pure water. The content of the tube was centrifuged for 5 min at 3,100 × g (4 °C) and the fibril-containing supernatant was collected. This extraction step was repeated for a total of ten times.

***Preparation of amyloid fibrils* in vitro**

Amyloid fibrils were prepared *in vitro* from a number of polypeptide chains that were either obtained through recombinant expression systems, custom chemical synthesis, commercial sources or purified from patient tissue (see Table 1 for details). The required amount of each fibril forming polypeptide chain was measured gravimetrically with a LA120S analytical balance (Sartorius) and dissolved in 1 mL of the respective solvent. The precise conditions of each fibrillation reaction are provided in Table 1. Except from the FOR005 LC fragment, we prepared agitated samples by placing the sample tubes onto an MTS 2/4 digital microtiter shaker (IKA) at 150 rpm.

***Purification of SAP***

500 µL from the first Tris–EDTA buffer fraction obtained during fibril extraction were mixed with 2 mL 7.5 M guanidine hydrochloride in 20 mM sodium phosphate buffer, pH 6.5, incubated overnight on a platform shaker (Polymax 1040, Heidolph) set to 200 rpm. Trifluoroacetic acid (TFA) was added to generate a TFA concentration in the sample of 0.2 % (v/v) and loaded onto a 3 mL Resource RPC column (GE Healthcare) that was equilibrated with solution A [0.1 % (v/v) TFA in water]. After loading the sample onto the column, the column was washed with solution A and the elution was initiated by applying a linear gradient of 0 to 100 % solution B [0.1 % (v/v) TFA in 86 % (v/v) acetonitrile]. Fractions of 3 mL were collected. SAP-containing fractions were pooled and dialysed two times for 1 hour each and once overnight against 1 L pure water using a 3.5 kDa Spectra/Por 6 dialysis membrane (Spectrum Laboratories). The pure SAP was lyophilized and reconstituted in pure water. 200 µL of this sample were mixed with 800 µL 7.5 M guanidine hydrochloride in 25 mM sodium phosphate buffer, pH 6.5, and the protein concentration was determined based on the extinction at 280 nm using the Lambert-Beer law. The theoretic extinction coefficient of SAP was 51,005 [$M^{-1}$ $cm^{-1}$]. The SAP solution was afterwards adjusted with pure water as required.

***Quantification of SAP in* ex vivo *fibril samples***

The SAP concentration of fibril extracts containing 200 µg/mL fibrils was determined by using semi-dry western blot. To that end, the fibril samples were initially diluted with pure water as described in the legend of Fig. S4. The samples were subjected to denaturing protein electrophoresis. The gel was transferred via semi-dry western blot onto a PVDF membrane at 20 V for 35 min in transfer buffer [1x NuPAGE Transfer Buffer, 20 % (v/v) methanol, Thermo Fischer]. Afterwards the membrane was washed in PBS-T [137 mM NaCl, 2.7 mM KCl, 10 mM $Na_2HPO_4$, 2 mM $KH_2PO_4$, 0.05 % (v/v) Tween 20, pH 7.4] for 5 min and blocked for 1 h in PBS-T containing 5 % (w/v) dry milk powder on a platform shaker (Polymax 1040, Heidolph). The membrane was then transferred into 20 mL PBS-T containing 5 % (w/v) dry milk powder and anti-SAP primary antibody (rabbit, Abcam, ab45151, 1:5,000, overnight, 4 °C under constant agitation). Afterwards the membrane was washed three times for 10 min with PBS-T and transferred into 20 mL PBS-T containing 5 % (w/v) dry milk powder and anti-rabbit secondary antibody coupled to horse radish peroxidase (goat, Dako, 1:2,000, 1 h, 4 °C under constant agitation). After three wash steps with PBS-T (10 min each), the membrane was treated with west pico plus chemiluminescent substrate solution (Thermo Fischer) and imaged with Box Chemi XL1.4. The SAP concentration was determined by densitometry by comparison with a standard (Fig. S4) in a fashion similar to the quantification of the extracted fibrils.

**SI Tables**

**SI Table 1 Conditions of *in vitro* fibril formation.**

The source of the polypeptide chain is indicated in the second column: R: recombinant; V: commercial vendor; C: chemical synthesis. The third column provides the concentration (Conc.) of the fibril forming polypeptide chain in the incubation reaction. Except from apomyoglobin, which had to be generated from holomyoglobin as described [4], all other polypeptide chains were used as obtained without further reprocessing.

| **Polypeptide chain, species** | **Origin** | **Conc.** | **Incubation conditions** |
|---|---|---|---|
| Apomyoglobin, horse | V, Sigma | 1 mg/mL | 50 mM sodium borate, pH 9.0, 65°C, 2 days |
| Aβ(1-40), human | R, in house [5] | 1 mg/mL | 50 mM sodium borate, pH 9.0, room temperature, 7 days |
| α-Crystallin, bovine | V, Sigma | 10 mg/mL | 1 M guanidine hydrochloride, 60°C, 1 day |
| Glucagon, human | R, V, Sigma | 1 mg/mL | 50 mM glycine/HCl buffer, pH 2.5, room temperature, with agitation, 2 days |
| Immunoglobulin LC FOR(005), variable domain | R, In house [6] | 50 μM | 20 mM sodium phosphate buffer, pH 6.5, 50 mM NaCl, 37°C, 120 rpm, 2 weeks |
| Insulin, bovine | V, Sigma | 1 mg/mL | Pure water, pH 2.0, 60°C, 3 days |

| | | | |
|---|---|---|---|
| Insulin, human | R, V, Sigma | 1 mg/mL | Pure water, pH 2.0, 60°C, 3 days |
| α-Lactalbumin, bovine | V, Sigma | 5 mg/mL | 100 mM NaCl, pH 2.0, 37°C, with agitation, 7 days |
| β-Lactoglobulin, bovine | V, Sigma | 4 mg/mL | Pure water, pH 2.0, 80°C, 7 days |
| Lysozyme, human | R, V, Sigma | 14 mg/mL | Pure water, pH 2.0, 65°C, 14 days |
| SAA1.1, human | R, in house [7] | 10 mg/mL | 50 mM $Na_2HPO_4$, room temperature, pH 1.0, 4 days |
| SAA1.1, murine | R, in house [8] | 5 mg/mL | 50 mM $Na_2HPO_4$, pH 3.0, 37°C, 4 days |
| SEVI, human | C, gift from J. Münch, Ulm University Clinic, Germany | 2.5 mg/mL | 137 mM NaCl, 2.7 mM KCl, 10 mM $Na_2HPO_4$, 2 mM $KH_2PO_4$, pH 7.4, 37°C, with agitation, 7 days |
| α-Synuclein, human | R, gift from K. Danzer, Ulm University Clinic, Germany | 1 mg/mL | 137 mM NaCl, 2.7 mM KCl, 10 mM $Na_2HPO_4$, 2 mM $KH_2PO_4$, pH 7.4, 37°C, with agitation, 3 days |
| TTR | R, gift from J. W. Kelly, Scripps Research Institute, U.S.A. | 2.5 mg/mL | Pure water, pH 2.0, 37°C, 3 days |

**SI Figures**

**SI Figure 1**

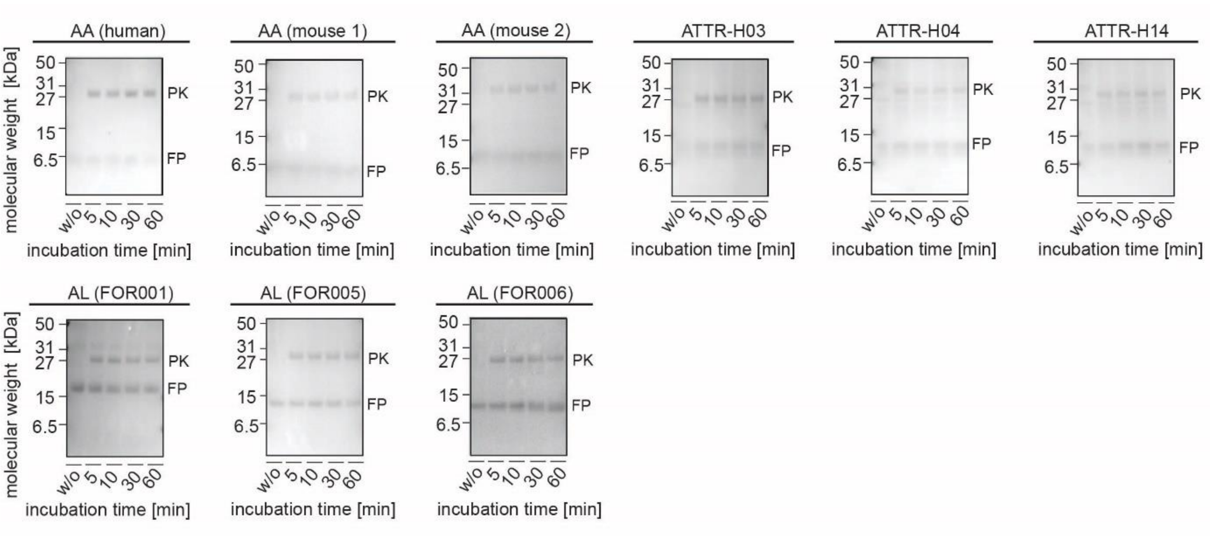


**SI Figure 1.**

***Ex vivo* amyloid fibrils are highly proteinase K stable.**

In contrast to Fig. 1 these samples contain a nominal fibril protein concentration of 100 µg/mL and 20 µg/mL proteinase K. Nine cases of *ex vivo* amyloid fibrils were examined. Each panel shows a Coomassie stained denaturing protein gel of the fibrils digested with proteinase K for 5, 10, 30 or 60 min, as indicated in the Figure. The first lane shows fibril sample before proteinase K addition. FP: fibril protein; PK: proteinase K.

**SI Figure 2**

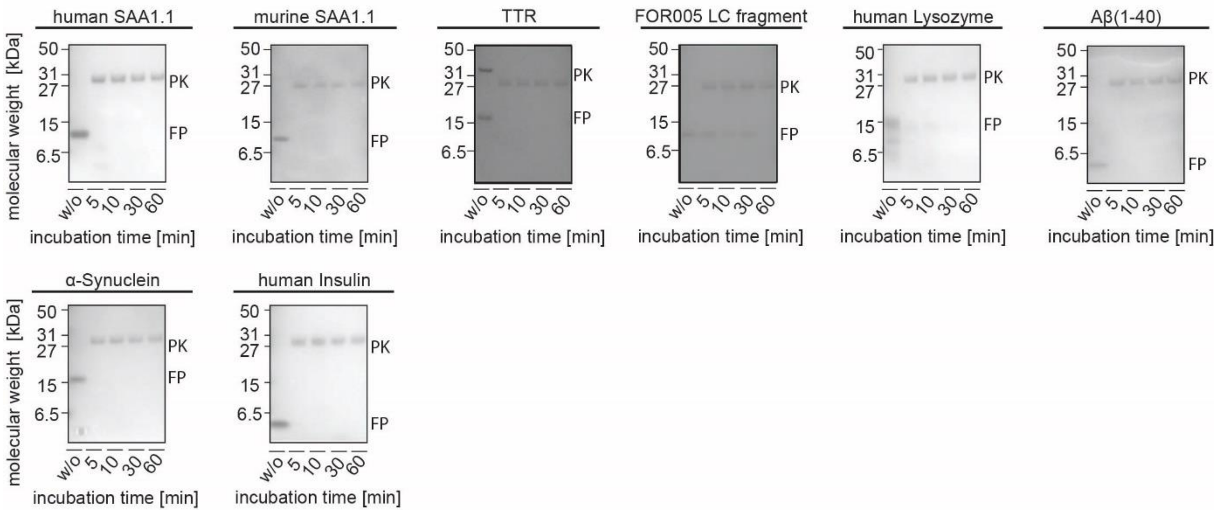


**SI Figure 2.**

***In vitro* fibrils from disease-associated polypeptide chains are proteinase K sensitive.**

In contrast to Fig. 2 these samples contain a nominal fibril protein concentration of 100 µg/mL and 20 µg/mL proteinase K. Eight cases of *in vitro* fibrils from disease-associated polypeptide chains were examined. Each panel shows a Coomassie stained denaturing protein gel of the fibrils digested with proteinase K for 5, 10, 30 or 60 min, as indicated in the Figure. The first lane shows fibril sample before proteinase K addition. FP: fibril protein; PK: proteinase K.

**SI Figure 3**

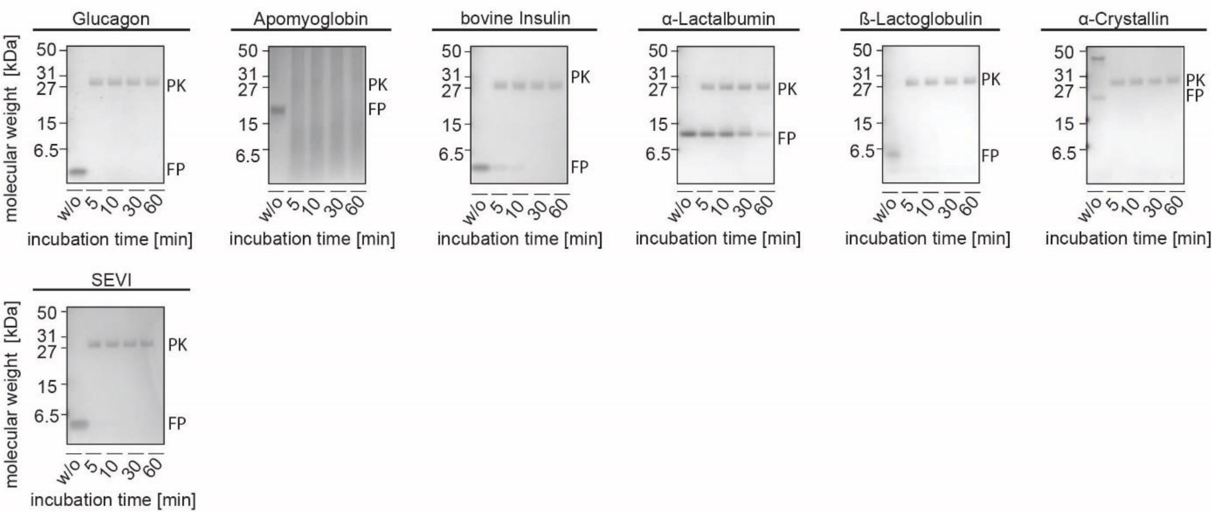


**SI Figure 3.**

**Most *in vitro* fibrils from non-pathogenic polypeptide chains are proteinase K sensitive.**

In contrast to Fig. 3 these samples contain a nominal fibril protein concentration of 100 µg/mL and 20 µg/mL proteinase K. Seven cases of *in vitro* fibrils from non-pathogenic polypeptide chains were examined. Each panel shows a Coomassie stained denaturing protein gel of the fibrils digested with proteinase K for 5, 10, 30 or 60 min, as indicated in the Figure. The first lane shows fibril sample before proteinase K addition. FP: fibril protein; PK: proteinase K.

**SI Figure 4**

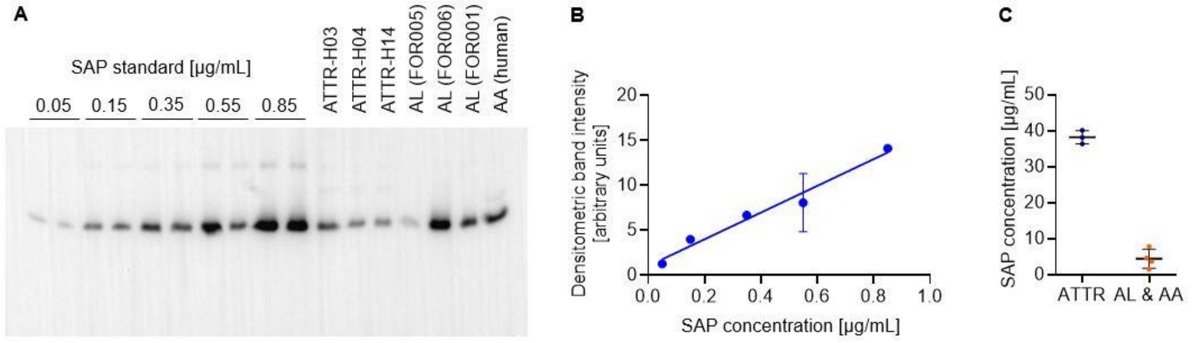


**SI Figure 4.**

**Quantification of SAP in the *ex vivo* fibril samples.**

(A) Western blot of the SAP standard and of different ATTR, AL and AA fibril samples. The analyzed fibril samples contained 200 µg/mL fibrils and were diluted 1:10 (AL and AA samples) or 1:100 (ATTR-H03) and 1:200 (ATTR-H04, ATTR-H14) prior tp preparation of the gel. (B) Densitometric quantification of the SAP standard (n = 2), fitted with a straight line. (C) Quantification of SAP in the seven fibril samples analyzed in panel (A) resulted in mean concentration values of 38.2 ± 1.5 µg/mL for the ATTR samples (n = 3) and 3.4 ± 2.3 for the AL and AA samples (n = 4). Errors represent standard deviation.
